# SpaDiff: Denoising for Sequence-based Spatial Transcriptomics via Diffusion Process

**DOI:** 10.1101/2025.10.07.681011

**Authors:** Jiazhang Cai, Yongkai Chen, Luyang Fang, Wenxuan Zhong, Guo-Cheng Yuan, Ping Ma

## Abstract

Spatial transcriptomics enables transcriptome-scale analysis with spatial resolution but suffers from spot-swapping, where RNA molecules drift from their true locations, introducing noise and reducing spatial specificity. We introduce SpaDiff, a denoising method that treats spot-swapping as a diffusion process. SpaDiff simulates the displacement of RNA molecules and reverses it to restore their original spatial distribution while preserving molecular counts. Evaluations on simulated and real data demonstrate that SpaDiff enhances spatial specificity of gene expression, improves data integrity, and supports more accurate downstream analyses such as clustering and spatial domain identification.

## Introduction

Spatial transcriptomics has emerged as a pivotal technology for mapping gene expression within intact tissue architecture, revealing fundamental mechanisms in brain function, cancer progression, and cellular differentiation processes^1–5^. Unlike single-cell RNA-seq, spatial transcriptomics preserves the in situ context, enabling direct analysis of cellular niches, tissue organization, and cell-cell communication^6^. Commercialization through platforms such as 10X Genomics have accelerated the widespread adoption of spatial transcriptomics, highlighting the need for high-quality data to maintain biological fidelity^7^.

However, there is a notable phenomenon associated with the sequencing-based spatial transcriptomics technology called spot-swapping^8^, where RNA molecules diffuse from their original locations to adjacent spots on a tissue slide (Figure 1 (A)). This phenomenon distorts spatial resolution, causing transcripts from distinct cell types to co-localize artificially and leading to erroneous interpretations of tissue organization and cellular interactions. Ideally, the probes in one spot only capture RNA released from the cells directly above the spot. However, due to the intrinsic diffusive nature of RNA molecules, transcripts often migrate across spots, making the captured RNAs not fully restricted to the tissue-covered areas. Empirical evidence confirms this phenomenon: for example, significant RNA UMI counts are routinely detected in uncovered regions of Visium slides, demonstrating non-localized migration. A particularly compelling demonstration of spot-swapping was presented by Ni et al.^8^, who used human–mouse chimeric tissue samples to reveal extensive cross-contamination between spatial spots. Their experiment showed that species-specific transcripts were detected in off-tissue or mismatched regions, providing clear evidence of RNA molecule diffusion across spot boundaries. This observation highlights the severity of the spot-swapping and motivates the need for effective computational denoising strategies to ensure biological fidelity. A more detailed analysis of this experiment is presented in the Results section.

**Figure 1:**
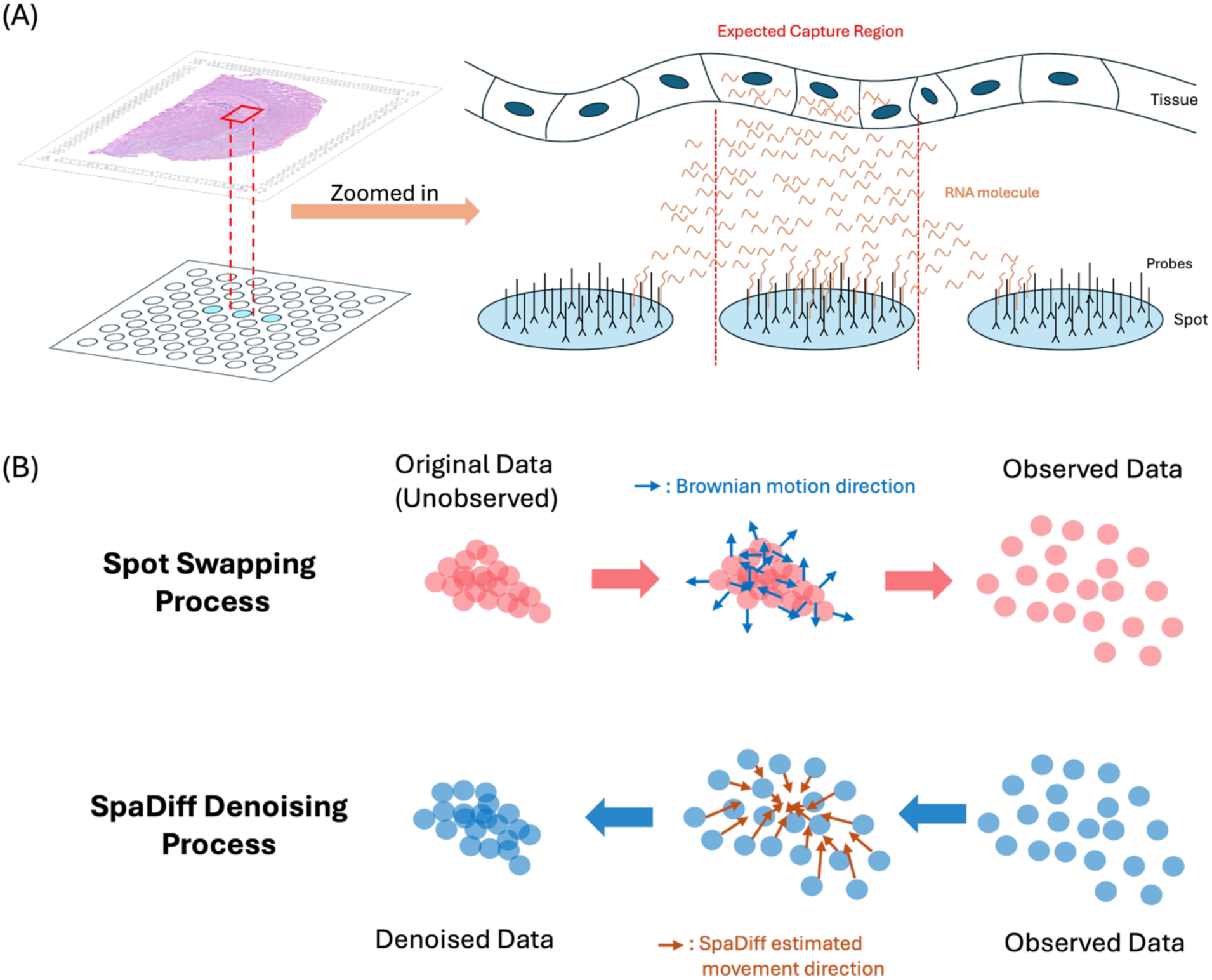
Introduction to the spot-swapping phenomenon and SpaDiff algorithm. (A) The illustration of the spot-swapping phenomenon. The RNA molecules from the expected capture region are captured not only by the spot under the region but also by its neighboring spots. (B) The illustration of SpaDiff’s reverse diffusion process correcting spot-swapping from observed data to noiseless data.

While several methods have been developed to denoise gene expression data^9–13^, such methods were originally designed for single-cell RNA-seq analysis, which does not contain spatial information. In contrast, methods that are specifically designed for spatial transcriptomics data are still lacking, and the few existing methods still have different limitations. SpotClean^8^ uses a probabilistic model to reconstruct the data based on the distribution after removing the effect of the spot-swapping phenomenon. It shows strong power in data denoising but fails to preserve the original RNA counts, potentially distorting biological insights. GF^14^, on the other hand, focuses on filtering out uniformly expressed genes to enhance spatially informative genes using optimal transportation, yet it does not effectively correct for RNA migration errors.

Diffusion models have recently gained attention for their effectiveness in denoising complex, structured data^15,16^. Unlike traditional smoothing methods that rely on fixed kernels, diffusion models adapt to data distributions^17–19^, making them well-suited for handling the non-Gaussian noise patterns commonly observed in biological datasets. In the context of spatial transcriptomics, spot-swapping inherently resembles a physical diffusion process, making diffusion models exceptionally well-suited for addressing this challenge. In this paper, we introduce SpaDiff, which leverages this analogy by utilizing the forward diffusion process to simulate the spot-swapping effect and then employing the reverse diffusion process for precise data denoising, which is illustrated in Figure 1 (B). By focusing on the movement of individual RNA molecules, SpaDiff offers a novel and elegant solution that preserves the biological integrity of the spatial transcriptomics data, enhancing both the accuracy and reliability of downstream analyses. In both synthetic and real datasets, including human brain and chimeric samples, SpaDiff outperforms existing methods in spatial specificity, clustering accuracy, and biomarker enrichment analysis.

## Results

### Overview of SpaDiff

SpaDiff is designed to mitigate the spot-swapping effect by employing a diffusion framework that mirrors the underlying physical process (Figure 1B). The forward diffusion step models how the true, underlying spatial gene expression is progressively corrupted by noise until it resembles the observed, noisy data. Conversely, the reverse process starts from the observed data and incrementally removes this noise to recover the underlying clean signal, serving as the denoising mechanism.

Unlike standard generative diffusion models, which start from pure white noise and generate new samples by reversing the entire process, SpaDiff only leverages the forward–reverse structure to describe the relationship between the ground truth and the observed data. This partial use of the diffusion framework allows SpaDiff to focus specifically on modeling and correcting spot-swapping noise, without requiring complex generative training or large datasets. Moreover, because spatial transcriptomics data inherently consist of two-dimensional spatial expression maps for each gene, the score function that guides the reverse process can be estimated directly using mathematical techniques. This makes SpaDiff particularly well-suited for efficient denoising in spatial transcriptomics.

SpaDiff models the spot-swapping phenomenon using Brownian motion, a stochastic process that captures the random movement of particles, which is analogous to the diffusion of RNA molecules across spatial spots. SpaDiff operates on the observed gene count matrix by iteratively adjusting the position of each RNA molecule based on a score function derived from each gene’s spatial distribution. In the model, the dynamics are decomposed into a stochastic noise term and a drift term, with the latter defined by the score function. The score function introduces directionality by guiding the motion toward regions consistent with the data distribution. During denoising, this drift effectively directs each RNA molecule toward regions of higher probability density, which are interpreted as the molecule’s true spatial origin.

Intuitively, SpaDiff calculates the gradient of the data distribution, where molecules are most likely to belong, and “pulls” each observed point back in that direction. This mechanism allows SpaDiff to eliminate the noise introduced by spot-swapping and recover the original spatial structure. This approach not only preserves the spatial specificity of the RNA data but also enhances the accuracy of subsequent biological interpretation. As illustrating examples, we consider two idealized scenarios where the spatial expression pattern of a gene can be represented by two canonical 2D shapes: a double moon and a double circle, respectively (Figures 2(A), (B)), whereas the observed data show similar patterns but the boundaries are significantly blurred due to the spot-swapping effect simulated by adding Gaussian noise (Figures 2 (C),(D)). These shapes are widely used in the machine learning literature to evaluate algorithms’ ability to recover nonlinear boundaries and topological structure under noise. Although not biologically derived, they provide contrasting geometries: double-moons test SpaDiff’s ability to denoise curved, non-convex manifolds, while double-circles examine its handling of radial symmetry and separation. Together, they serve to demonstrate the model’s ability to recover diverse spatial patterns without requiring prior biological assumptions.

**Figure 2:**
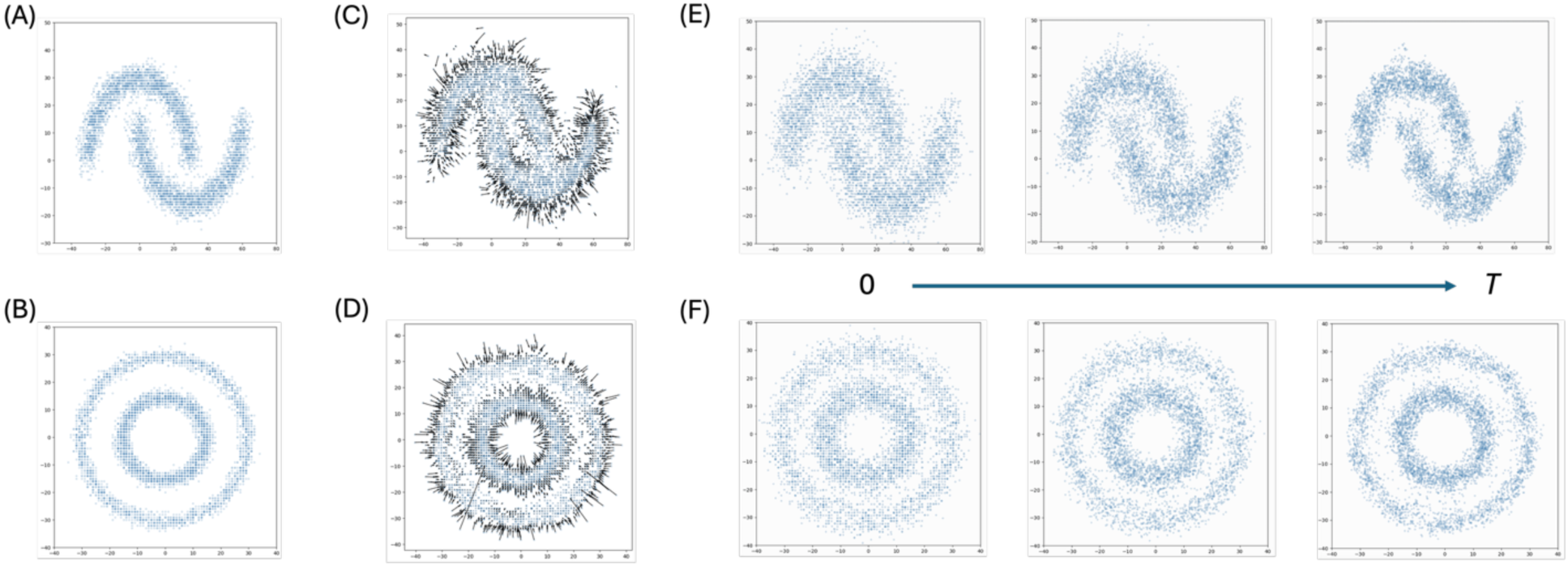
Conceptual demonstration of SpaDiff on synthetic data. Two synthetic examples are used to visually demonstrate how SpaDiff processes spatial data. (A-B) Ground truth synthetic datasets used as the reference. (C-D) Noisy data obtained by adding noise to the ground truth, with corresponding estimated score functions. (E-F) Denoising process of SpaDiff, showing the transformation from noisy observations to denoised spatial patterns that largely reproduce the ground truth.

SpaDiff computes the score function for the overall distribution of each gene, which essentially acts as the scaled gradient directing each point toward the center of its distribution. The magnitude of the score function reflects the intensity of the restoring force that drives points back toward their distribution center. This process mirrors the reversal of the spot-swapping effect. As shown in Figures 2 (E), (F), SpaDiff progressively denoises the data using its learned score function. These examples visually demonstrate SpaDiff’s ability to estimate spatial density gradients and guide noisy points back toward high-probability regions.

### Simulation of Spot-Level Spatial Transcriptomics Data

To evaluate the performance of our denoising method for spot-level spatial transcriptomics, we developed a simulation framework that generates realistic sequence-based data using high-resolution single-molecule transcriptomic measurements from a publicly available 10x Xenium mouse brain dataset as ground truth^20^. Conceptually, this process can be visualized as placing a slide with an array of capture spots onto the original single-molecule data to mimic how Visium spots capture transcripts within a defined physical area, as shown in Figure 3 (A). Starting from each detected UMI’s gene identity and precise spatial coordinates, we introduce local transcript diffusion by applying a Brownian motion perturbation to each molecule’s position to mimic the phenomenon of spot swapping observed in sequence-based platforms such as Visium, where transcripts may diffuse across neighboring capture areas. By controlling the standard deviation of this Brownian motion, we can systematically vary the degree of mixing among spots, allowing us to compare the performance of our denoising method under different levels of spatial signal blurring. To replicate the physical layout of a Visium slide, we construct a hexagonal grid of capture spots with a diameter and center-to-center spacing matching the platform’s specifications, typically around 55 µm in diameter and 100 µm apart^21^. For each spot, we aggregate all UMIs whose Brownian-perturbed coordinates fall within the spot’s circular capture area, ignoring those that fall into the inter-spot gaps, as occurs in real experiments. By design, this simulation framework provides a clear and controllable link between the blurred, spot-level observation and the high-resolution ground truth, enabling rigorous benchmarking of denoising and deblurring methods under conditions that closely mimic the biological and technical variability of real Visium experiments, as shown in Figure 3 (B).

**Figure 3:**
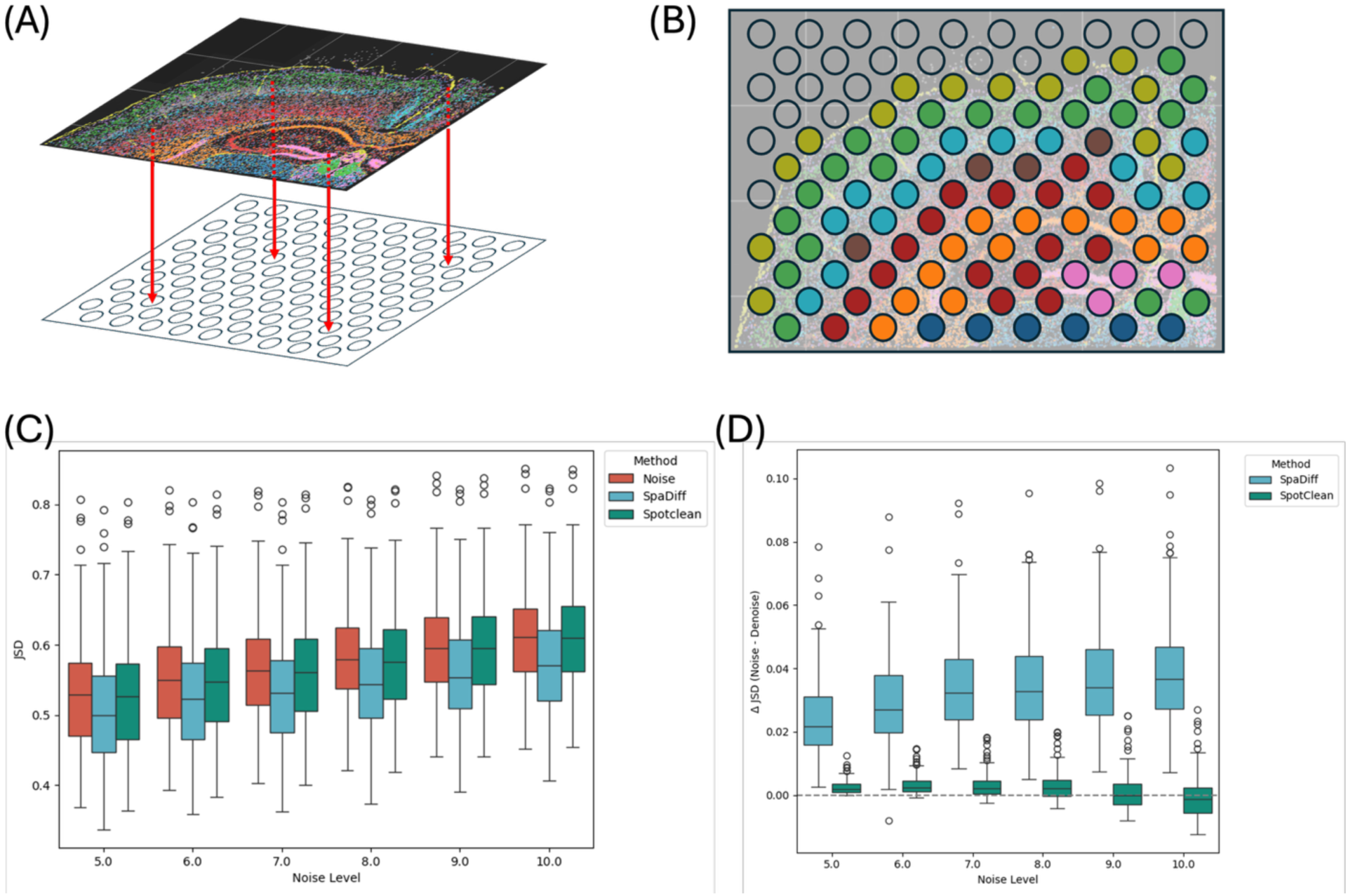
Evaluation of SpaDiff on synthetic data generated from Xenium data. (A) Conceptual illustration of simulating Visium spot-level measurements by capturing Xenium UMIs within spatially defined hexagonal spots. (B) Illustration of the synthetic Visium data. (C) Comparison of Jensen–Shannon divergence (JSD) between the noisy data and ground truth versus the denoised data, across varying noise levels. (D) Per-gene comparison of JSD reduction after denoising using SpaDiff and SpotClean, highlighting the improved recovery of spatial structure by SpaDiff.

To quantitatively evaluate denoising performance, we compared the outputs of SpaDiff with the original noisy data and an existing method, SpotClean, using Jensen–Shannon divergence (JSD)^22^ as a measure of similarity to the ground truth. For each simulated dataset, we computed gene-wise JSD between the noisy spot-level observations and the original high-resolution Xenium-derived reference, then repeated the same calculation after applying denoising. To assess the robustness of each method under varying levels of spatial diffusion, we repeated this evaluation across a range of Brownian motion noise levels. This allowed us to test performance under increasingly severe spot mixing.

Figure 3 (C) shows that, when aggregating the JSD across genes, SpaDiff consistently showed lower divergence from the ground truth compared to the noisy input, although the reduction was modest and not statistically significant across all conditions. In contrast, SpotClean failed to show a consistent improvement and often produced distributions of JSD that were indistinguishable from or slightly worse than the unprocessed noisy data.

To further investigate method performance on a per-gene level, we computed the ΔJSD for each gene, defined as the difference in JSD before and after denoising. Figure 3 (D) shows that SpaDiff exhibited a significant shift in ΔJSD toward positive values across all noise levels, indicating a consistent improvement in spatial fidelity after denoising. In contrast, SpotClean showed ΔJSD distributions centered around zero, suggesting little to no benefit when applied to single-gene denoising in this context.

These results highlight the effectiveness of SpaDiff in recovering spatial signals from noisy observations, particularly in cases where individual genes exhibit strong and localized spatial patterns. Furthermore, the use of high-resolution ground truth allows us to interpret these improvements with confidence, validating our simulation framework as a rigorous tool for benchmarking spatial denoising methods.

### Performance on the DLPFC tissue

To evaluate SpaDiff’s performance in a complex real-world setting, we applied it to a widely used sequence-based spatial transcriptomics dataset: the 10x Genomics Visium human dorsolateral prefrontal cortex (DLPFC) dataset from the LIBD study^23^. This dataset consists of 12 tissue sections collected from three neurotypical adult donors.

Each section was manually annotated into seven cytoarchitecturally defined regions, including six cortical layers (Layers 1 through 6) and white matter (WM), based on histological features and known gene expression markers. One annotated slice is shown in Figure 4 (A).

**Figure 4:**
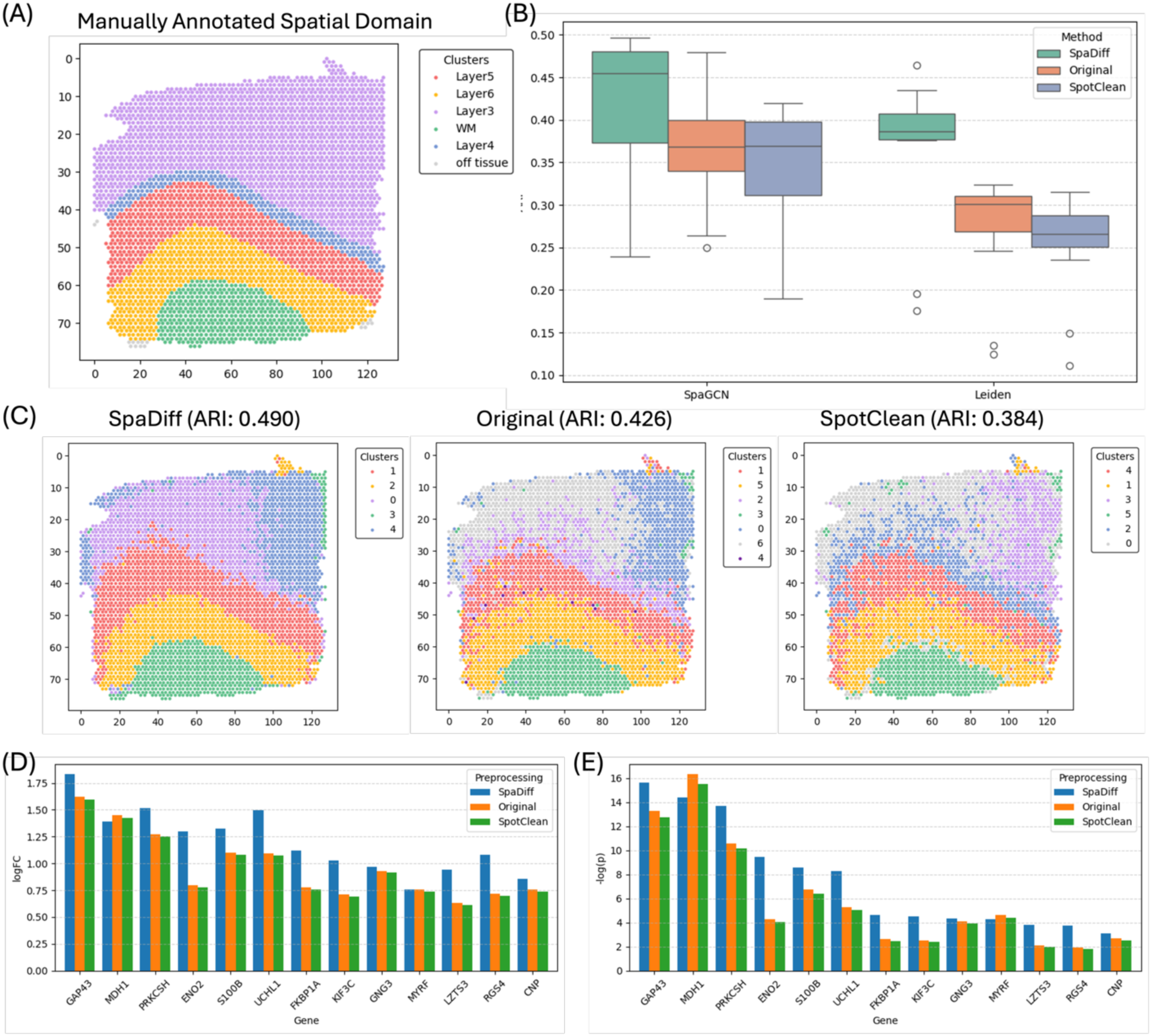
Evaluation of the performance of SpaDiff on DLPFC datasets. (A) Manual annotation of one DLPFC tissue. Boxplot of the ARI between the clustering result of Leiden and SpaGCN using the original data and the denoised data from SpaDiff and Spotclean, and the manual annotation on all twelve tissues. (C) Domain detection result using SpaGCN on the original data and the denoised data using Spadiff and SpotClean, respectively. (D) Comparison of log fold change between white matter and layer 6 for some biomarker genes. (E) Comparison of the negative log p-value for the same genes.

We assessed denoising effectiveness using two clustering strategies: Leiden^24^ and SpaGCN^25^. Leiden, a widely used graph-based clustering algorithm, does not incorporate spatial information and thus serves as a lower-bound benchmark. In contrast, SpaGCN is a spatially aware method that integrates gene expression with spatial context. We select SpaGCN as one representative example of existing spatial methods. This allows us to test whether SpaDiff can enhance downstream analyses regardless of the specific algorithm used. By including both a non-spatial baseline (Leiden) and a spatially informed approach (SpaGCN), we show that SpaDiff can improve results in both settings.

For each clustering method, we compared results from the original data, SpaDiff-denoised data, and SpotClean-denoised data against manual annotations using the Adjusted Rand Index (ARI). As shown in Figure 4 (B), SpaDiff consistently achieved higher ARI scores than both the raw and SpotClean-denoised datasets under both Leiden and SpaGCN. For Leiden, the improvement from SpaDiff was substantial and highly significant. For SpaGCN, SpaDiff still outperformed the other two inputs, but the improvement compared to Leiden was smaller than the other two. This is because SpaDiff-denoised data already exhibited a relatively clear and well-defined spatial pattern, allowing even Leiden, which does not incorporate any spatial information, to achieve performance approaching that of SpaGCN.

Interestingly, SpotClean sometimes performed worse than the raw data, particularly in tissues with higher technical noise. This may be attributed to its reliance on a parametric probabilistic model, which can be sensitive to model misspecification or data quality. In contrast, SpaDiff’s nonparametric diffusion-based framework imposes fewer assumptions on the data-generating process, allowing it to perform robustly across diverse conditions, including noisy or low-quality tissue sections such as those found in the DLPFC dataset.

To further evaluate biological interpretability, we examined domain detection using SpaGCN on a representative tissue section (Figure 4 (C)). In SpaDiff-denoised data, spatial domains appeared more distinct and coherent, aligning more closely with manual annotations than either raw or SpotClean-denoised data.

We also quantified the ability to distinguish spatial regions by comparing the log fold change and statistical significance of known biomarker genes between WM and Layer 6^23^ (Figures 4 (D) and 4 (E)). Across samples, SpaDiff-denoised data exhibited larger log fold changes and smaller p-values than both raw and SpotClean-denoised data, demonstrating its ability to amplify biologically meaningful differences between regions.

Overall, these results indicate that SpaDiff improves the quality of spatial transcriptomics data in a way that benefits a range of downstream analyses, from basic clustering without spatial information to spatially aware methods, while also producing data so well-structured that even non-spatial approaches can approach the performance of spatially informed methods.

### Performance on the human-mouse chimeric samples

To evaluate the effectiveness of SpaDiff in correcting spatial transcriptomics noise under controlled conditions, we tested the model on human–mouse chimeric tissue samples from a study by Ni et al.^8^. This dataset presents a unique experimental setup where human and mouse tissues are intentionally mixed within the same spatial transcriptomics slide. As shown in Figure 5 (A), the dataset provides experimentally derived ground truth annotations, clearly delineating human-only, mouse-only, and mixed tissue regions. Unlike most spatial transcriptomics datasets, which rely on indirect or manual annotation based on histology and marker gene expression, this dataset enables direct and objective validation of denoising performance. It is particularly valuable for assessing spot-swapping artifacts, as UMI molecules originating in one species that are captured in the other provide clear indicators of cross-contamination.

**Figure 5:**
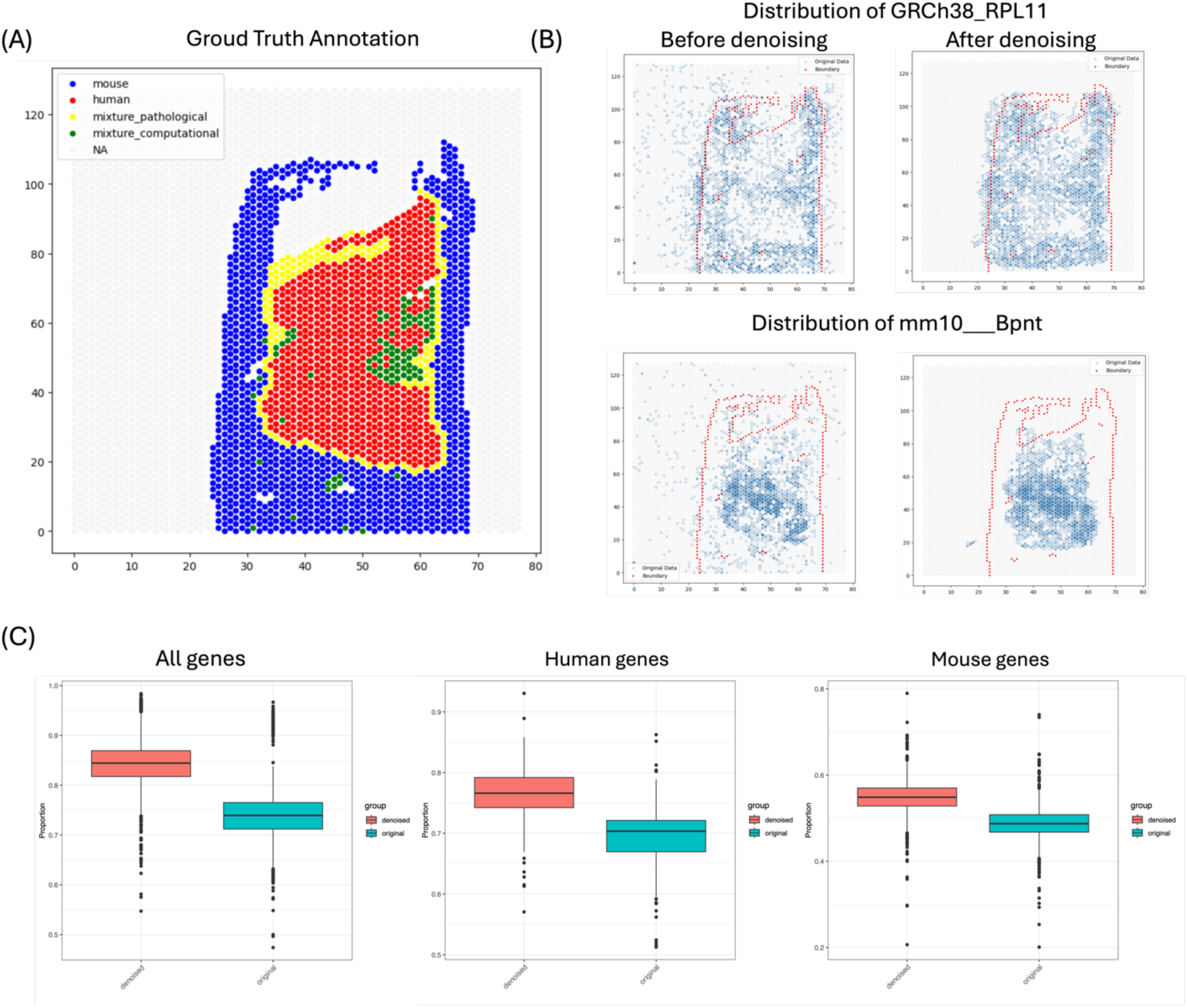
Performance of SpaDiff on human-mouse chimera tissue. (A) Ground truth annotation of the human and mouse tissue area. (B) Distribution of one human and mouse marker genes before and after denoising using SpaDiff. Comparison of the proportion of reads in-tissue area for all genes, the proportion of reads in the human-tissue area for all human genes, and the proportion of reads in the mouse-tissue area for all mouse genes before and after denoising using SpaDiff.

In conventional preprocessing pipelines, UMI counts detected outside the tissue-covered regions are often excluded under the assumption that they represent noise. However, recent work^8^ has shown that a non-negligible portion of off-tissue UMI counts may originate from the tissue itself, likely due to spot-swapping or diffusion effects.

Although downstream analyses are often restricted to in-tissue spots for interpretability, recent evidence shows that many off-tissue UMI counts originate from tissue due to spot-swapping. Discarding these signals risks losing genuine biological information and can introduce systematic bias, especially if spot-swapping affects marker gene localization. By explicitly modeling and correcting for this diffusion, SpaDiff preserves signal integrity across the full slide, improving downstream accuracy.

Figure 5 (B) visualizes SpaDiff’s denoising performance using two canonical marker genes: GRCh38_RPL11 (human) and mm10_Bpnt (mouse). The two plots on the left show their spatial distribution in the raw data. Prior to denoising, both genes are detected beyond their expected regions: human gene reads are present in mouse tissue areas, and mouse gene reads appear in human regions, strongly confirming the presence of spot-swapping noise. The right-hand plots in Figure 5 (B) show the gene distributions after applying SpaDiff. The denoised expression maps reveal a striking improvement in spatial specificity: the mouse gene becomes restricted to peripheral areas corresponding to mouse tissue, while the human gene is concentrated more centrally, in line with the ground truth annotation. These results visually confirm SpaDiff’s ability to relocalize transcripts to their true spatial origin.

To evaluate performance quantitatively, we assessed the proportion of gene counts localized to their appropriate tissue compartments. As shown in Figure 5 (C) (left), we calculated, for each gene, the fraction of reads that fall within the actual tissue area (as opposed to background or cross-species regions). After denoising with SpaDiff, this proportion increases significantly, indicating improved signal-to-noise ratio and biological coherence. We further stratified the analysis by species: the middle and right boxplots in Figure 5 (C) show the proportion of human gene reads in human tissue and mouse gene reads in mouse tissue, respectively. Across both categories, SpaDiff consistently improves alignment between gene origin and spatial location, providing strong evidence of effective spot-swapping correction.

Notably, this chimeric dataset provides a rare opportunity to evaluate denoising performance against a known spatial ground truth. However, many alternative denoising approaches rely on pre-filtering spots outside of tissue regions at the outset, which would preclude assessment of spot-swapping noise in precisely these areas. As such, benchmarking SpaDiff against these methods in this context is infeasible and would not reflect their intended use. By contrast, SpaDiff directly models and corrects cross-spot contamination without discarding spatial information, enabling robust recovery of biological signal even in highly artifact-prone regions.

In summary, the human–mouse chimeric dataset offers a rare opportunity to validate denoising methods against known spatial ground truth. SpaDiff not only resolves cross-species contamination at the level of individual genes but also improves overall data structure across the transcriptome. These results demonstrate that SpaDiff can restore biological fidelity in noisy spatial transcriptomics datasets and is well-suited for preprocessing data in complex or artifact-prone tissue settings.

## Discussion

Sequence-based spatial transcriptomics is pivotal for comprehensive gene profiling, leveraging high-throughput sequencing technologies to detect gene expression with genome-wide coverage and high molecular specificity. However, these techniques are often limited by technical artifacts—most notably the spot-swapping phenomenon, in which mRNA molecules diffuse and are captured by neighboring barcoded spots. This artifact introduces substantial noise, distorts the true spatial distribution of gene expression, and negatively impacts downstream analyses such as clustering, spatial domain identification, and biomarker detection.

SpaDiff addresses this critical challenge using a principled approach based on diffusion modeling, which mimics the reverse dynamics of spot swapping to restore molecular localization. Importantly, the method adheres to physical constraints: the total number of UMI counts in the denoised data is exactly preserved, maintaining the integrity and interpretability of quantitative gene expression. This conservation property is particularly crucial when SpaDiff is used as a preprocessing step, ensuring compatibility with existing downstream pipelines. Moreover, SpaDiff employs a unified generative model to denoise the entire dataset, treating it as an interconnected system in which molecular movements are governed by consistent probabilistic rules. This cohesive modeling strategy not only simplifies implementation but also enhances the method’s robustness and theoretical soundness.

The efficacy of SpaDiff has been demonstrated through applications to both synthetic and real-world datasets. Empirical results show that SpaDiff not only corrects spot-swapping artifacts but also improves the biological plausibility of spatial gene expression patterns. After denoising, key biomarker genes exhibit refined spatial localization, and aberrant UMIs captured outside of tissue regions are appropriately reassigned. These improvements directly benefit downstream analyses, enabling more accurate spatial domain delineation, cell type localization, and biomarker discovery.

Looking ahead, SpaDiff opens several avenues for future development. Its modular and scalable architecture makes it compatible with larger and more complex datasets, including multi-section or 3D spatial transcriptomics. Furthermore, its continuous modeling approach may facilitate integration with histological imaging and cross-modal alignment, offering a promising bridge between sequencing-based and imaging-based spatial transcriptomics.

In summary, SpaDiff provides a powerful, principled solution to key challenges in sequence-based spatial transcriptomics. By simultaneously addressing spot-swapping artifacts, enhancing spatial resolution, and supporting scalable analysis, SpaDiff represents a significant advancement in preprocessing technology and paves the way for more accurate and insightful spatial transcriptomic studies.

## Conclusion

This study establishes SpaDiff as a robust denoising framework that corrects spot-swapping artifacts in spatial transcriptomics while preserving molecular counts. By enhancing the spatial fidelity of gene expression, SpaDiff improves biomarker detection, spatial domain identification, and overall data integrity. Its generalizable modeling strategy and compatibility with existing pipelines underscore its importance as a practical preprocessing step. Beyond current applications, SpaDiff sets the stage for scalable, multimodal spatial analyses, representing a significant step toward more accurate and comprehensive interpretation of spatially resolved transcriptomic data.

## Methods

### Algorithm overview

The core mechanism of SpaDiff is the application of the diffusion model to achieve the goal of denoising the data. Suppose the expression level of one gene on the spot from a sequence-based spatial transcriptomics data is denoted by 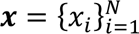, where *N* is the number of spots. The spot swapping phenomenan and its reverse process can be modeled using a typical diffusion model with a forward process to add noise and a reverse process to remove the noise. Mathematically, we can use the stochastic differential equation (SDE) to represent the round-trip processes:

Forward process:

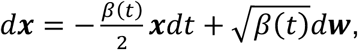

Reverse process:

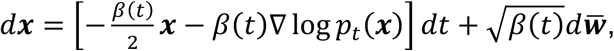

where *t* represents the time, β(*t*) is the noisy scale, usually in the form of a pre-defined monotonously increasing function of *t*, ***w*** is a standard Brownian motion, ***w̅*** is the reverse-time Brownian motion, which is also a standard Brownian motion, but appears only in the reverse process, ∇ log *p_t_*(*x*) is the score function, which is calculated by the gradient of the logarithm of the probability density function (PDF) of data *x* at time *t*. Under our assumption, the spot-swapping follows a basic diffusion process, where the diffusion is only caused by the Brownian motion of the molecules. Therefore, we can remove the drift term and simplify the two SDEs to:

Forward process:

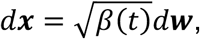

Reverse process:

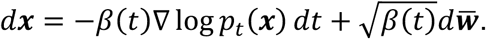

Further, under our assumption, we can define σ_*t*_ = β(*t*) = β*t*, which is a linear function of time *t*. In this case, σ_*t*_ is actually the variance of the noise. The discrete form of the forward and reverse process can be expressed as:

Forward process:

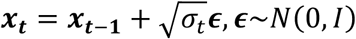

Reverse process:

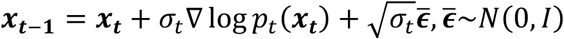

It should be noted that the reverse process is to infer *x*_*t*–1_ given *x*_*t*_. Therefore, the sign before the score function in the discrete form is different from the one in the SDE form. With the discrete formula, we assume the observed data *x*_*T*_ is diffused from the hidden clean data *x*_0_ after *T* steps.

In the forward and reverse processes of the diffusion model, estimating the score function ∇ log *p*_*t*_(*x*_*t*_) becomes challenging if the distribution *p*_*t*_(*x*_*t*_) lacks a closed-form expression or is difficult to approximate. This issue commonly arises when working with image or text data. In such cases, the diffusion model typically treats the score function as a function of data *x*_*t*_ and time *t*, and employs a deep neural network to train it. However, under our assumption, the score function can be directly estimated in a more efficient and accurate way. In SpaDiff, each gene’s denoising process is approached as a simplified two-dimensional scatter data challenge. Unlike the complex deep learning-based score function estimation methods, SpaDiff employs a kernel-based method, which is well-developed to estimate the score function^26^ for a straightforward, mathematically explicit, and faster computation of the score function. Specifically, we approximate the score function *s_p_*(*x*) = ∇ log *p*(*x*) within a reproducing kernel Hilbert space (RKHS) using a curl-free kernel, which ensures the estimated score function is a valid gradient field. The score function is obtained by minimizing a regularized Fisher divergence. By applying integration by parts under appropriate boundary conditions, the loss function can be reformulated into a formula that depends only on the gradient of the estimated score function *s*(*x*), eliminating the need to explicitly know *s_p_*(*x*), which results in the final objective:

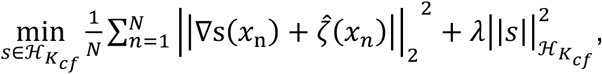

where *N* is the number of observation, ℋ_*K_cf_*_ represents the reproducing kernel Hilbert space (RKHS) with a curl-free kernel, 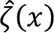 is an empirical estimator of the divergence of the score function, replacing the unknown true gradient of the score function, and ensures that the optimization problem remains solvable using only the given samples. There are two norms in the above expression: the first one is the ℒ_)_ norm, and the second one is the norm in the Reproducing Kernel Hilbert Space, measuring the smoothness of the function. To make the algorithm more efficient, we implement the *v*-method^27^, an iterative approximation technique applying a polynomial approximation of the inverse operator, to avoid direct matrix inversion, which can be computationally expensive and unstable.

Denote the estimated score function given data *x* as *s̃*(*x*), then we can define the denoise model of SpaDiff as follows:

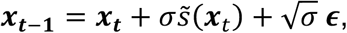

where *t* is the time step, ranging from 0 to *T*, *x*_*t*_ represents the data at time *t*, *ε̃N*(0, *I*) is a standard Brownian motion, and σ is a constant representing the variance of the noise. SpaDiff treats the initially observed data as *x*_*T*_ and denoises the data step by step until the denoised data *x*_0_.

SpaDiff contains two tuning parameters, which flexibly adapt to different scenarios. One is the variance of the noise, which highly depends on the platforms and the sample preprocessing. The other is the kernel width during the score function calculation, which is highly reliant on the shape of the tissue sample. We set the default parameters of SpaDiff to a noise variance of 0.1 and a kernel width of 15.0. As SpaDiff includes a coordinate rescaling step prior to denoising, these default settings are broadly applicable across datasets without requiring dataset-specific tuning. In practice, we find that this configuration consistently yields robust performance across a wide range of scenarios.

## Algorithm acceleration

To further enhance the effectiveness and scalability of the denoising process, we introduce four strategies to the core SpaDiff algorithm. Each addresses a key challenge in balancing denoising quality, computational efficiency, and scalability for large spatial transcriptomics datasets.

1. Density-Aware Score Function Scaling: During score function estimation, we observe that values tend to be larger in densely populated regions and lower in sparser areas. However, effective denoising requires the movement of UMI molecules from low-density to high-density regions, which contradicts the default directionality of the score function. To resolve this, SpaDiff introduces a density-aware scaling factor that adjusts the magnitude of the score function based on local point density. Specifically, the scaler is inversely proportional to the local density: points in sparsely populated regions receive a higher scaling factor, increasing the score magnitude and pushing them toward denser, more biologically plausible areas. This adjustment ensures that the denoising trajectory is aligned with the underlying spatial patterns and enhances the biological interpretability of the reconstructed gene expression fields.
2. Subsampled Score Estimation for Computational Efficiency: The gradient-based score function depends on all observed data points, making it computationally expensive, especially for large-scale spatial transcriptomics datasets. However, since the score is defined on the global distribution, estimating it using the entire dataset is often unnecessary. To improve computational efficiency, SpaDiff employs a random subsampling strategy to approximate the score function using only a subset of data points. Each individual point’s score is then evaluated using this approximation. This significantly reduces computation time while preserving the quality of the denoising, making the method more scalable for real-world applications. It can also be easily extended to other subsampling strategies, which have been proven to be powerful in different scenarios^28–30^.
3. Drift Scaling for Accelerated Convergence: In theory, the denoising process, formulated as the reverse of a stochastic diffusion process, is guaranteed to converge when the step size is infinitesimally small. However, such small steps result in a large number of iterations, leading to a considerable computational burden in practice. To accelerate convergence, we introduce a drift-scaling parameter that increases the magnitude of the drift term. This modification effectively pushes the distribution toward its high-density regions more aggressively, reducing the number of steps needed while maintaining a close approximation of the ideal reverse process. The drift-scaling parameter strikes the balance between theoretical precision and computational feasibility, allowing SpaDiff to produce high-quality denoised outputs in significantly less time. In practice, we find that moderate drift scaling accelerates denoising with minimal impact on final accuracy, making it a valuable heuristic for performance tuning in large datasets.
4. Parallel Gene-wise Denoising for Scalability: SpaDiff models each gene independently, making the denoising process parallelizable across genes. It enables the use of multi-core or distributed computation, significantly reducing wall-clock time for large datasets. This gene-wise parallelism ensures that SpaDiff scales well even with datasets containing tens of thousands of genes and thousands of spatial locations. By exploiting this structure, SpaDiff maintains both high performance and practical usability for large-scale spatial transcriptomics studies.

## Declarations

### Ethics approval and consent to participate

Not applicable.

### Consent for publication

Not applicable.

### Availability of data and materials

The code for SpaDiff has been uploaded to GitHub at: https://github.com/JiazhangCai/SpaDiff. The data used in this study include both simulated and real datasets and are all public available datasets. Simulation data were generated using high-resolution Xenium mouse brain datam which can be found at: https://www.10xgenomics.com/datasets/xenium-prime-fresh-frozen-mouse-brain. For real data, we analyzed the human dorsolateral prefrontal cortex (DLPFC) dataset available from the SpatialLIBD project, available at: http://research.libd.org/spatialLIBD/, as well as the mouse–human chimeric tissue dataset generated with the 10x Genomics Visium platform, available at: https://www.ncbi.nlm.nih.gov/geo/query/acc.cgi?acc=GSE178221.

### Authors’ contributions

Conceptualization, G.-C.Y., P.M., W.Z., J.C., and Y.C.; methodology, P.M., Y.C., and J.C.; coding, L. F. and J.C.; formal analysis, Y.C., J.C, F.L., G.-C.Y., and P.M.; writing – original draft, J.C.; writing – review & editing, J.C., G.-C.Y., L.F., P.M, Y.C., and W.Z.; supervision, G.-C.Y., P.M., and W.Z.; funding acquisition, G.-C.Y., P.M., and W.Z.

## Acknowledgements

Not applicable.

